# Idioblasts as pathways for distributing water absorbed by leaf surfaces to the mesophyll in *Capparis odoratissima*

**DOI:** 10.1101/2019.12.17.879577

**Authors:** Juan M. Losada, Miriam Díaz, N. Michele Holbrook

## Abstract

*Capparis odoratissima* is a tree species native to semi-arid environments of the northern coast of South America where low soil water availability coexists with frequent nighttime fog. Previous work with this species demonstrated that *C. odoratissima* is able to use water absorbed through its leaves at night to enhance leaf hydration, photosynthesis, and growth.

Here, we combine detailed anatomical evaluations of the leaves of *C. odoratissima,* with water and dye uptake experiments in the laboratory. We used immunolocalization of pectin and arabinogalactan protein epitopes to characterize the chemistry of foliar water uptake pathways.

The abaxial surfaces of *C. odoratissima* leaves are covered with overlapping, multicellular peltate hairs, while the adaxial surfaces are glabrous but with star-shaped “structures” at regular intervals. Despite these differences in anatomy, both surfaces are able to absorb condensed water, but this ability is most significant on the upper surface. Rates of evaporative water loss from the upper surface, however, are coincident with cuticle conductance. Numerous idioblasts connect the adaxial leaf surface and the adaxial peltate hairs, which contain hygroscopic substances such as arabinogalactan proteins and pectins.

The highly specialized anatomy of the leaves of *C odoratissima* fulfills the dual function of avoiding excessive water loss due to evaporation, while maintaining the ability to absorb liquid water. Cell-wall related hygroscopic compounds present in the peltate hairs and idioblasts create a network of microchannels that maintain leaf hydration and promote the uptake of aerial water.

## INTRODUCTION

Water-energy dynamics intrinsically drive global patterns of plant diversity (Kreft and Jetz, 2007), but the global increase of aridity will likely make plants more vulnerable to the lack of water in the soil (Choat et al., 2018; Olson et al., 2018). In arid and semiarid environments, the traditional soil-atmosphere continuum concept of hydraulic redistribution in plants (Nobel, 2009), is compromised due to prolonged soil drought. Xerophyllous plants thrive in these ecosystems thanks to leaf anatomical adaptations that reduce rates of water loss, such as smaller areas, lower stomata index and impermeable coating structures, like cuticles and trichomes (see Shields, 1950). Surprisingly, some of the structures that prevent evaporative water loss may also facilitate aerial water uptake into the leaves, decoupling soil water uptake from that of the canopy (Benzing et al., 1978; Gouvra and Grammatikopoulos, 2003).

Foliar water uptake (FWU) is a widespread phenomenon of vascular plants, known for three centuries, and empirically tested in at least 53 plant families (Dawson and Goldsmith, 2018). FWU has direct consequences on plant function, relaxing tension in the water column of the xylem, enhancing turgor-driven growth, and influencing the hydraulic dynamics of agro and ecosystems (Mayr et al., 2014; Steppe et al., 2018; Aguirre-Gutiérrrez et al., 2019). The key conditions for FWU are met in fog-dominated environments (Tognetti, 2015; Weathers et al., 2019), where high atmospheric humidity enhances nighttime dew formation on leaf surfaces, thus increasing the possibility of foliar water absorption. Indeed, studies in montane cloud forests of Brazil (Eller et al., 2016), coastal California redwood forests in the USA (Burgess and Dawson, 2004), or cloud forests in Mexico (Gotsch et al., 2014), have demonstrated that aerial water is an important input contributing to plant function, especially during the dry periods of the year. In contrast, foliar uptake in semiarid areas of the tropics is less studied, despite being a pivotal water source that enhances biomass productivity (Díaz and Granadillo, 2005; Limm et al., 2009). Better characterization of the pathways of foliar absorption will enhance understanding of the mechanisms semiarid plants use to hydrate leaves with aerial water.

Some species absorb water through the natural leaf openings, such as stomata (Burkhardt et al., 2012; Berry et al., 2014) or hydathodes (Martin and von Willert, 2000). Stomatal water uptake alone conflicts with CO_2_ dynamics between leaves and the atmosphere (Boanares et al., 2019). Other structures that are supposedly impermeable to water further participate in the aerial water uptake, including cuticles (Vaadia and Waisel 1963; Yates and Hutley,. 1995; Fernández et al., 2017; Schuster et al., 2017), and trichomes (Franke 1967; Benzing et al. 1978; reviewed by Berry et al., 2019). While the use of these wall-thickened, sealing structures appears as a good strategy to capture atmospheric water condensed on the leaf surfaces, demonstration of their hygroscopic capacity requires a careful examination of their cell wall biochemistry, so far lacking for most plant species. Early works in this line showed that the absorptive capacity of leaves relates with the presence of polysaccharides under the cuticle (Kerstiens, 1996). However, the role of ubiquitous compounds of the cell walls, such as pectins and glycoproteins, on foliar water uptake, has been scarcely reported (Boanares et al., 2018). The question remains on whether sealing structures such as trichomes or sclerenchymatous structures displayed such hygroscopic compounds.

This gap of knowledge affects particularly to xerophyllous species, which display unique anatomies with abundant sclerenchymatous tissues, such as idioblasts, highly specialized structures that are understudied from a functional perspective. Idioblasts (Schwendener, 1874) are thick walled cells typically buried in the mesophyll of leaves of vascular plants (Bailey and Nast, 1945; Tomlinson and Fisher, 2005), and recurrently found in the leaves of xerophyllous species (Foster, 1956; Heide Jorgensen, 1990). They have been traditionally explored from the perspective of morphology, ontogeny, and taxonomic value (Foster,1944; 1945a,b; Bloch, 1946; Foster, 1955a,b; Rao and Mody, 1961). However, important functions adscribed to idioblasts are merely inferred from their putative stiffness, such as support and defense (Foster, 1947; Tucker, 1964; Rao and Sharma, 1968), or from their topology, such as a possible role in leaf capacitance (Heide-Jorgensen, 1990). Some noteworthy exceptions suggest that idioblasts could affect the physiology of the leaves more than previously thought, such as serving as light guides (Karabourniotis, 1998). The diverse array of anatomies of xerophyllous species is exemplified in the genus *Capparis* (Rao and Mody, 1961; Gan et al., 2013). *Capparis odoratissima* is a species native to the semiarid tropical environments of the South American continent with a remarkable capacity to produce new biomass using canopy irrigation as the only water source (Díaz and Granadillo, 2005), likely through foliar water uptake. However, the mechanisms of this uptake are unknown.

Here we focus on *C. odoratissima* with the goal of understanding the relationship between anatomy and function in xerophylous leaves. We show that the leaves of *C. odoratissima* are highly specialized structures that perform a dual function: first, they are armed organs that minimize water loss, but they further contain an intricate network of hygroscopic pathways that enhances s water uptake, thus maintaining leaf hydration upon water condensation on the leaf surface.

## MATERIALS AND METHODS

### Plant material

Five stems with a minimum of 10 leaves of *Capparis odoratissima* were shipped in a plastic bag with a wet paper tissue from the University of Francisco de Miranda at the state of Falcón (Venezuela), and three days later received for evaluation in the laboratory of the Arnold Arboretum of Harvard University in Boston (MA, USA).

### Evaluation of the external leaf anatomy

Five to six fully expanded leaves were first scanned with a scale using a CANON CanoScan LiDE 400 scanner, and leaf areas were calculated with the Image J1.51d software (National Institutes of Health, Bethesda, MD, USA). These same leaves were then used to photograph the details of both adaxial and abaxial surfaces with a Zeiss Discovery v12 Dissecting microscope (objective 0.63x PanApo). Adaxially, the idioblast tips were easily visualized by their translucency, and, abaxially, the peltate hairs were counted using their central parts as references. Both idioblasts and peltate hairs were counted in 1mm^2^ areas at five different positions in each leaf surface. To count the number of stomata, five leaves were fixed in FAA (formalin:acetic acid, Johansen 1940), washed three times in distilled water, one hour each, stained with Feulgen reaction (modified from Barell and Grossniklaus, 2005). They were finally cleared with a solution containing ethanol:benzoyl benzoate 3:1 (v/v) for six hours, ethanol:benzoyl benzoate 1:3 (v/v), and finally benzoyl benzoate: dibutyl pftalate 1:1 (v/v) for several days (Crane and Carman, 1987). Following removal of the peltate hairs, 1mm^2^ samples with guard cells stained in red, were imaged with a Zeiss LSM700 Confocal Microscope connected to a AxioCam 512 camera and the Zen Blue 2.3 software, objective was 20x/0.8 M27 Plan-Apochromat. Note that the area occupied by the rounded base of the peltate hairs had not stomata in the abaxial epidermis (3.45%), and hence it was subtracted from the total area of leaves to get a more accurate picture of stomatal frequency.

### Evaluation of the internal leaf anatomy: idioblast characterization

To better understand the structure of the idioblasts in the mesophyll, hand transverse thin sections (i.e. perpendicular to the leaf surface) of the individual leaves were obtained and stained with a solution of 0.01% w/v calcofluor white in 10mM CHES buffer with 100mM KCl (pH=10) (Hughes & McCully, 1975), that stains cellulose, and then with a uramine-O in 0.05 M Tris/HCl buffer, (pH=7.2), was used to detect cutinized lipids (Heslop-Harrison and Shivanna, 1977). These sections were visualized with a Zeiss Axioskop microscope equipped with epifluorescence, and imaged with an AxioCam HRc camera, attached to the Zen Blue software with multichannel. For calcofluor white, the filter used was DAPI narrow (Zeiss 48702) excitation G 365, bandpass 12 nm; dichroic mirror FT 395; barrier filter LP397; for auramine O, the filter used was AF488/FITC/GFP (Zeiss Filter set 9) excitation 450-490; dichroic mirror 510; emission filter LP515. Then, we isolated the idioblasts from other leaf tissues by macerating 1mm^2^ leaf pieces in a solution containing acetic acid: hydrogen peroxide 1:1 (v/v) at 60° for 2 days. The debris of these samples was then mounted in distilled water or glycerol, and visualized with a Zeiss LSM700 Confocal Microscope. Rendering images were obtained without staining with the Objective: 63x/1.40 Oil DIC M27 Plan-Apochromat.

Samples obtained from five previously fixed leaves in FAA were washed in distilled water three times, 1h each, and dehydrated with an increasing gradient of aqueous ethanol concentrations (10%, 30%, 50%, 70%, 100% (x3)). They were then soaked in the infiltration solution of Technovit7100 (Electron Microscopy Sciences, Hetfield, PA, USA) for several weeks, and polymerized under high nitrogen concentration conditions. Blocks with samples oriented in a paradermal way (i.e. parallel to the leaf surface), were serially sectioned at 4µm in with a Leica RM2155 rotary microtome (Leica Microsystems, Wetlar, Germany). Sections were then stained with an aqueous solution of 0.025% toluidine blue for general structure of the leaf tissues (Feder and O’Brien, 1968), and finally imaged with a Zeiss AxioImager A2 Upright microscope microscope equipped with a AxioCam 512 camera and a Zen Blue Pro with multichannel software.

### Water and dye uptake experiments

We first monitored cuticle transpiration in five full expanded leaves by letting them dry with their petiole covered with parafilm at room temperature (average 23±3°C), and 30±3% of relative humidity (RH), weighting them every hour with a four digit precision scale. We then adjusted the weight loss to a linear regression function (*r*^2^=0.98; slope = −0.0012). After that, we calculated the relative leaf water content (RWC) using the formula RWC=[(FW_t_-DW)/(DW-FW_0_)], were FW_t_ is the fresh weight at time t, DW is the dry weight, and FW_0_ is the fresh weight at time 0 (Fig. 5A). We further calculated the initial water content of the leaves (Wi) as the difference between the fresh weight minus the dry weight by leaf unit area.

Five more leaves were weighed (Wt_0_) before being submerged in distilled water with the petiole sealed -but not submerged- for 15min (*wet cycle*). To eliminate the surface water, these leaves were centrifuged at 1800rpm for 2min using a Sorvall RC6 Ultracentrifuge, and immediately weighed afterwards with a precision scale (Wt_1_). Leaves were left to dry at room temperature for 15min (*dry cycle*), and weighed again before immersion (Wt_2_). The cycles were repeated five times in a row. Based on these data, and assuming that cuticle evapotranspiration is 0 when leaves are submerged, we first calculated the net water absorbed after the *wet cycle* as to:

1) ABSt_1_= (Wt_1_-Wt_0_)/ L_A,_

where ABSt_1_ is the net water absorbed at time t (*wet cycle,* uneven numbers), and L_A_ is leaf area in cm^2^.

After that, and considering similar cuticle transpiration in both leaf sides, the expected cuticle evapotranspiration (C_EVT_) for the *dry cycle* was calculated as to:

2) C_EVT_ = [(−0.0012 * t)*2]+ Wt_0_, where t is time in hours.

3) ABSt_*_ = (Wt_*_ - W_t*-1_ + C_EVT_)/ L_A,_

where ABSt_*_ is the calculated water absorbed after the *dry cycles* (at even numbers), and W_t*_ the weight of the leaves after the *dry cycles*. We then evaluated the observed water balances for each alternate period as to:

4) ΔABS = (ABS_t_+ ABS_t-1_) for each alternate period.

The calculated water lost during the dry cycles corresponded with the observed values, suggesting that weight loss due to cuticle conductance was so low that it did not offset the water absorbed by leaves. Thus, we then plotted the net water absorbed only during the *wet cycles*, and adjusted it to a linear regression function (*r*^2^=0.96; slope = 0.073) (Fig. 5B), revealing a linear water uptake when leaves are submerged.

The question remained on whether each leaf side contributed equally to water uptake. To understand it, five leaves were loaded with 350uL of distilled water evenly distributed in 35 droplets (10uL each) on the adaxial surface, and in five more leaves loaded on the abaxial surface. When the droplets disappeared from the sufaces, the leaves were weighted with a precision scale, loaded again with the same amount of water, and this cycle repeated for a total period of 9 hours. While the atmospheric humidity was around 30%, it is worth noting that we replicated the experiment at high humidity (70%) within a chamber, but the water droplet disappearance from the leaf surfaces was so slow that made the experiment impracticable in a continuous fashion. However, it suggested that cuticle conductance is much lower at natural conditions, which display atmospheric humidities ranging from 50-70% (Díaz and Granadillo, 2005). We then calculated the water gains as described below for the dry cycles, but assuming cuticle transpiration in only one side of the leaves [C_EVT_ = (−0.0012* t) + Wt_0_], and finally adjusted the results to a linear regression curve. For the adaxial surface, *r*^2^=0.81; slope = 0.0012 (Fig. 5C).

To understand the pathways of water uptake, we used a 1% aqueous solution of the apoplastic fluorescent dye tracer Lucifer Yellow (LY; CH dilithium salt; Sigma). 10uL droplets of the dye were applied in either the adaxial or the abaxial surfaces within a humidified chamber, and waited until they disappeared from the surface. A paper tissue was then used to wipe the traces of the dye from the leaf surface prior to obtaining transverse sections with a double edge razor blade. Sections were mounted in an aqueous solution of 70% glycerol, and immediately observed with a Zeiss LSM700 Confocal Microscope (wavelength 488nm). Similar sections of the same leaves, but in areas without the dye were used as negative controls.

### Immunolocalization of pectin and arabinogalactan glycoprotein epitopes

A preliminary test with the general dye Ruthenium red was used in freshly dissected leaves to evaluate the presence of neutral pectins (Colombo and Rascio, 1977). Similarly, a preliminary test for arabinogalactan protein (AGP) presence was applied to cleared leaves with a 2% solution of the chemical reagent beta glucosyl Yariv reagent in 0.15M NaC1 (Yariv, 1967). The β-GlcYR reacts with arabinogalactan proteins giving a red color upon precipitation.

To detect the presence of pectic and AGP epitopes in the leaves of *Capparis odoratissima*, two monoclonal antibodies that detect highly sterified pectins (JIM5), and AGPs (JIM8) (Carbosource services, Georgia, USA), were used. 4µm cross and paradermal sections of embedded leaves were incubated with 1XPBS thrice for five minutes each, then in 5% bovine albumin serum (BSA), then in 1XPBS five minutes, and with the undiluted primary monoclonal antibody for 45 min. They were washed three times in 1XPBS, and finally incubated again with an anti-Rat secondary antibody Alexa 488 conjugated with a fluorescent compound Fluorescein Isiothianate. Three final washes with 1XPBS preceded observations with a confocal microscope (see details below). Negative controls were treated in the same way, but substituting the primary antibody by a solution of 1%BSA in PBS.

## RESULTS

### Extrinsic leaf anatomy of *Capparis odoratissima*

Detailed evaluations of the oblong, astomatous leaves of *C. odoratissima* showed a dark green color in the adaxial surface (Fig. 1A). Numerous translucent spots were located in concave areas of the surface, which corresponded with the tips of the idioblasts (Fig. 1B). Idioblast tips interspersed between the epidermal cells and, after staining and clearing the tissue, displayed a star-like shape when observed from above (Fig. C). At higher magnification, each idioblast projected a narrow pore toward the adaxial surface of the leaf (Fig. 1D). Abaxially, the leaves showed a pale green color (Fig. 1E), which resulted from the total coverage of the lamina by an imbricated carpet of peltate hairs (Fig. 1F). These non-glandular hairs had variable size, but a uniform multicellular, umbrella-like base, with thick-walled filiform cells (Fig. 1G), and a central shield on top that protrudes to the exterior. Despite having very thick walls, this central part revealed symplasmic connections between its cells (Fig. 1H).

**Figure 1.**
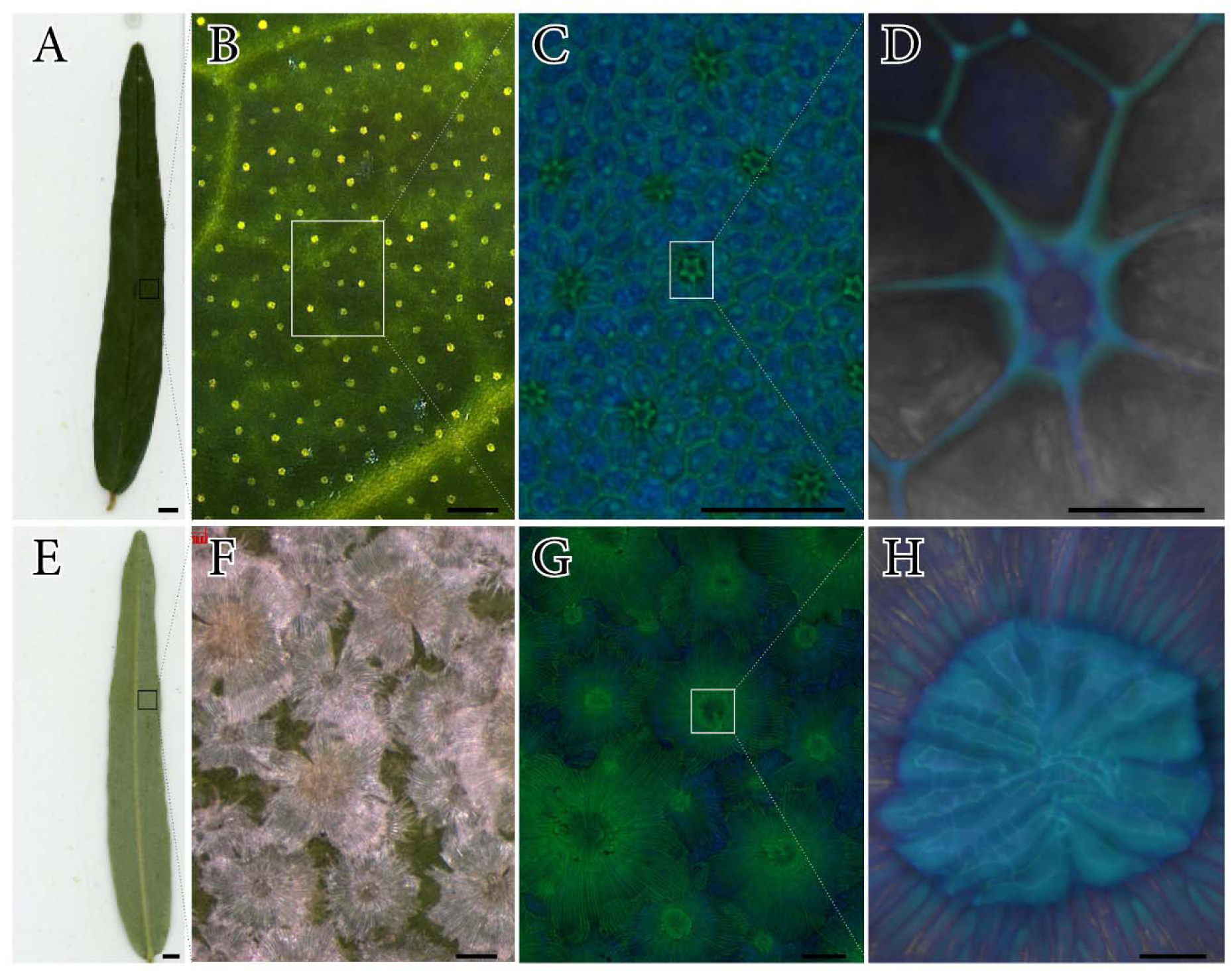
Leaf surface anatomy of *Capparis odoratissima*. **A** Elongated leaf shape with a bright green adaxial color. **B** Conspicuous pattern of translucent idioblast tips through the entire adaxial leaf area. **C** Cleared tissue showing a star-like appearance at the tip of these structures. **D** Closer view, showing a narrow pore in the center of the idioblasts. **E** Abaxial surface of the leaves with a pale green color. **F**. A continuous layer of imbricated peltate hairs covers the abaxial surface. **G** Peltate hairs are composed of two different units: the bigger one is underneath, with filiform projections that elongate laterally, and have thicker walls towards the center. **H**. In the center of each peltate trichome, a compact cup of thick walled filiform structures showed ample lumens. Dissecting scope images (A-D, adaxial; E-H, abaxial), and confocal images of cleared leaves stained with the Feulgen reaction (C,D, adaxial; G,H, abaxial).Scale bars: A,E = 500µm; B,F = 200µm; C,G = 100µm; D,H = 20µm.

### Intrinsic leaf anatomy of *Capparis odoratissima*: cuticles, peltate hairs and idioblasts

Cross-sections of the leaves revealed that the idioblasts are thick-walled columnar structures that connect the adaxial and the abaxial surfaces of the leaves (Fig. 2A, drawn by Metcalfe and Chalk, 1950). Idioblasts did not stain for lipids, but the auramine O staining revealed a thick cuticle layer in the adaxial surfaces of the leaves (Fig. 2B), which was thinner in the concave cavities on top of the idioblasts. In contrast, the thick columnar walls of the idioblasts stained intensely for cellulose with calcofluor white (Fig. 2C), which further delivered partial lignification upon phluoroglucinol staining. The abaxial surface of the leaf showed a more compact rugose cuticle under the peltate hairs (Fig. 2D). At the anchoring areas of the peltate hairs, the abaxial epidermis invaginated and contained a hyper accumulation of cutin (Fig. 2E).

**Figure 2.**
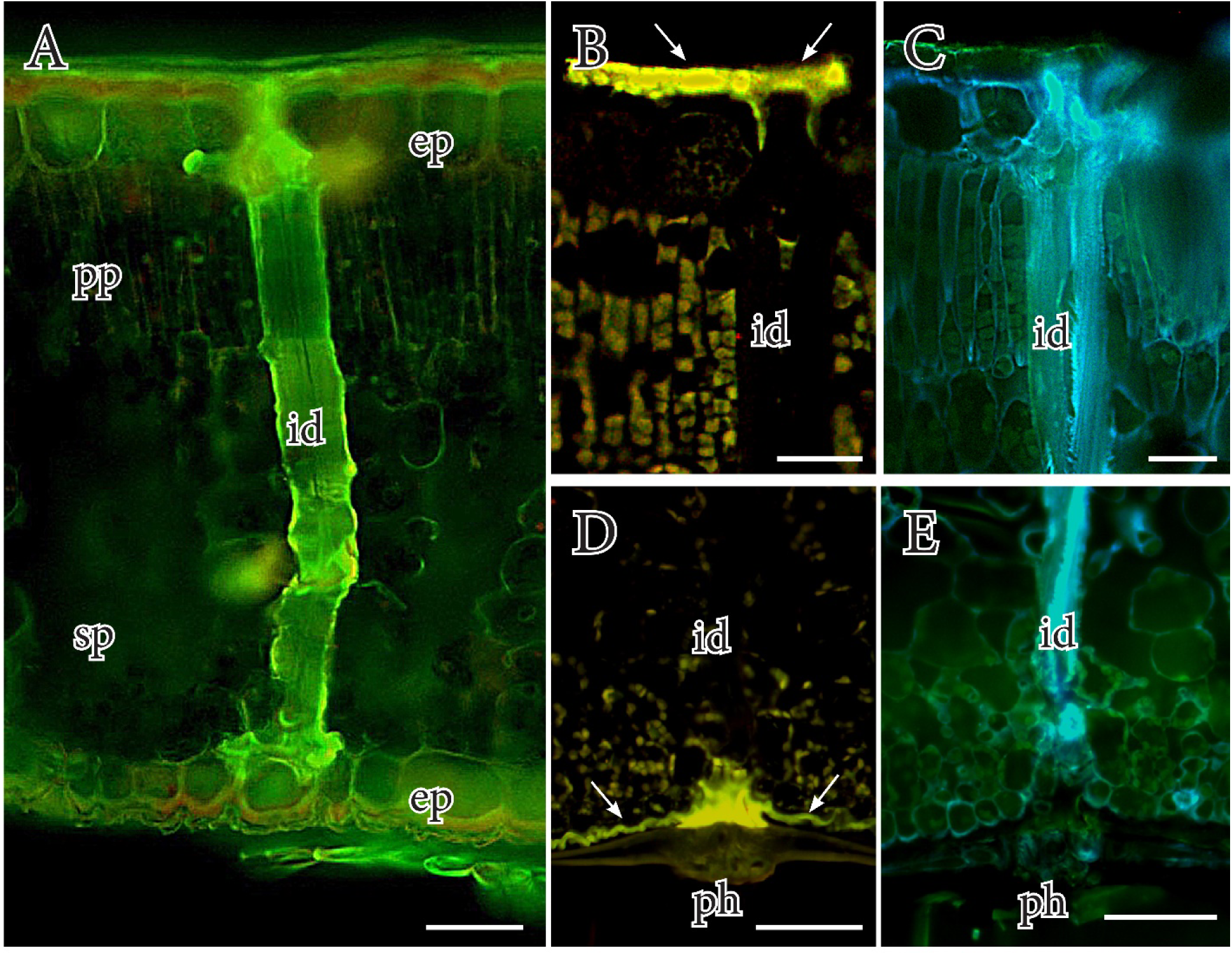
The internal anatomy of the leaves in *Capparis odoratissima*. **A** Cross-section of the leaf displaying one of the multiple ididoblasts that traverse the mesophyll and connects the adaxial and abaxial leaf surfaces. **B** Thick cuticle layer (white arrowheads) on the adaxial epidermis of the leaf, but thinner in the idioblas tips. **C** The thick walls of the idioblasts contain massive cellulose (blue color). **D** The cuticle in the abaxial surface (arrows) is thinner, rugose and accumulates massively at the base of the peltate hairs. **E** The idioblasts directly connect with the center of the peltate hairs in the abaxial surface of *C. odoratissima* leaves. Ep, epidermis; pp, palisade parenchyma; id, idioblast; sp, spongy parenchyma; ph, peltate hair. Scale bars: A,D,E = 50µm; B,C = 20µm.

Idioblasts exposed a sophisticated structure with a clear bipolar pattern from the adaxial to the abaxial side of the leaf (Fig. 3A). A thin projection that protruded apoplastically between cells of the upper epidermis connected with the surface, and was sustained underneath with four to five filiform anchors attached to the base of the epidermal cells (Fig. 3B). In the area of the palisade parenchyma, the idioblasts had a columnar shape, with a very thick wall, a narrow lumen, and crenations that connected the lumen with the palisade parenchyma (Fig. 3C). In the spongy mesophyll, the irregular shape of idioblasts was characterized by numerous protuberances and crenations that connected the lumen of the idioblasts with the mesophyll cells (Fig. 3D). While idioblasts thinned down and were irregularly shaped in the abaxial side of the leaf, they anastomosed and converged toward the base of the peltate hairs, yet keeping narrow lumens that connected the lumens of the idioblasts with the lumens of the peltate hairs (Fig. 3E). The average rate of idioblast anastomosing by peltate hair was 4:1 (Fig. 4), whereas the frequency of abaxial stomata under the peltate hairs was significantly lower.

**Figure 3.**
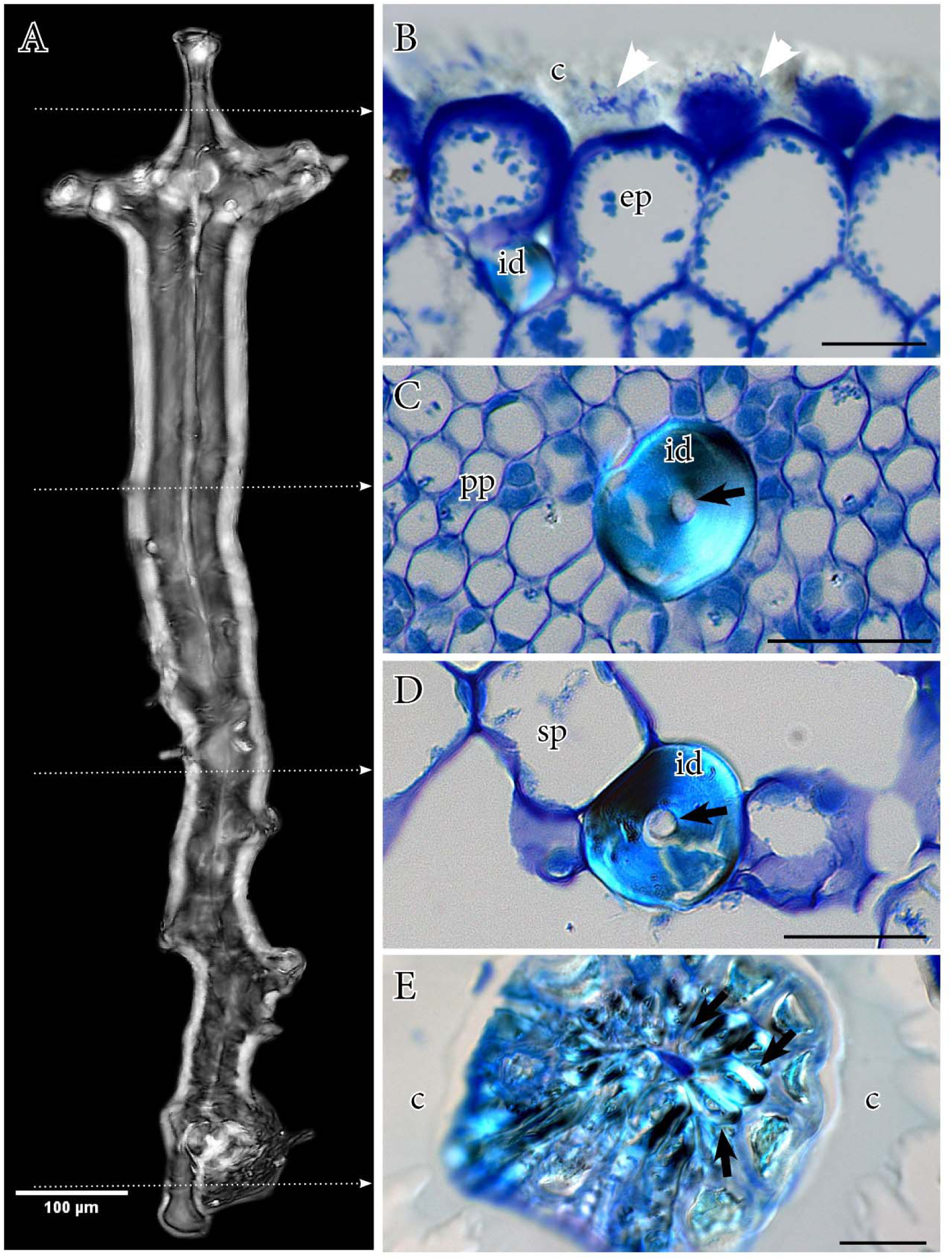
Characterization of the idioblasts of *Capparis odoratissima*. **A** Maximum 3D projection of an isolated idioblast showing their typical structure from a small tubular projection towards the adaxial epidermis (top) and the irregular walls at the bottom. **B** Cells of the adaxial epidermis are thick and have substantial extracellular materials that intersperse with the thick adaxial cuticle (white arrowheads). **C** The idioblasts are columnar at the area of the palisade parenchyma, with thick refringent walls and narrow lumens (arrow). **D** At the level of the spongy mesophyll, numerous canals connect the lumen of the idioblast with the cells of the leaf. **E** At the abaxial epidermis, adjacent idioblasts anastomose and form an irregular structure that coincides with the center of the peltate hairs. C, cuticle; ep, epidermis; pp, palisade parenchyma; id, idioblast; sp, spongy parenchyma; ph, peltate hair. Dotted arrows in A indicate the plane of section of images on the right. Scale bars: A = 100µm; B-E = 20µm.

**Figure 4.**
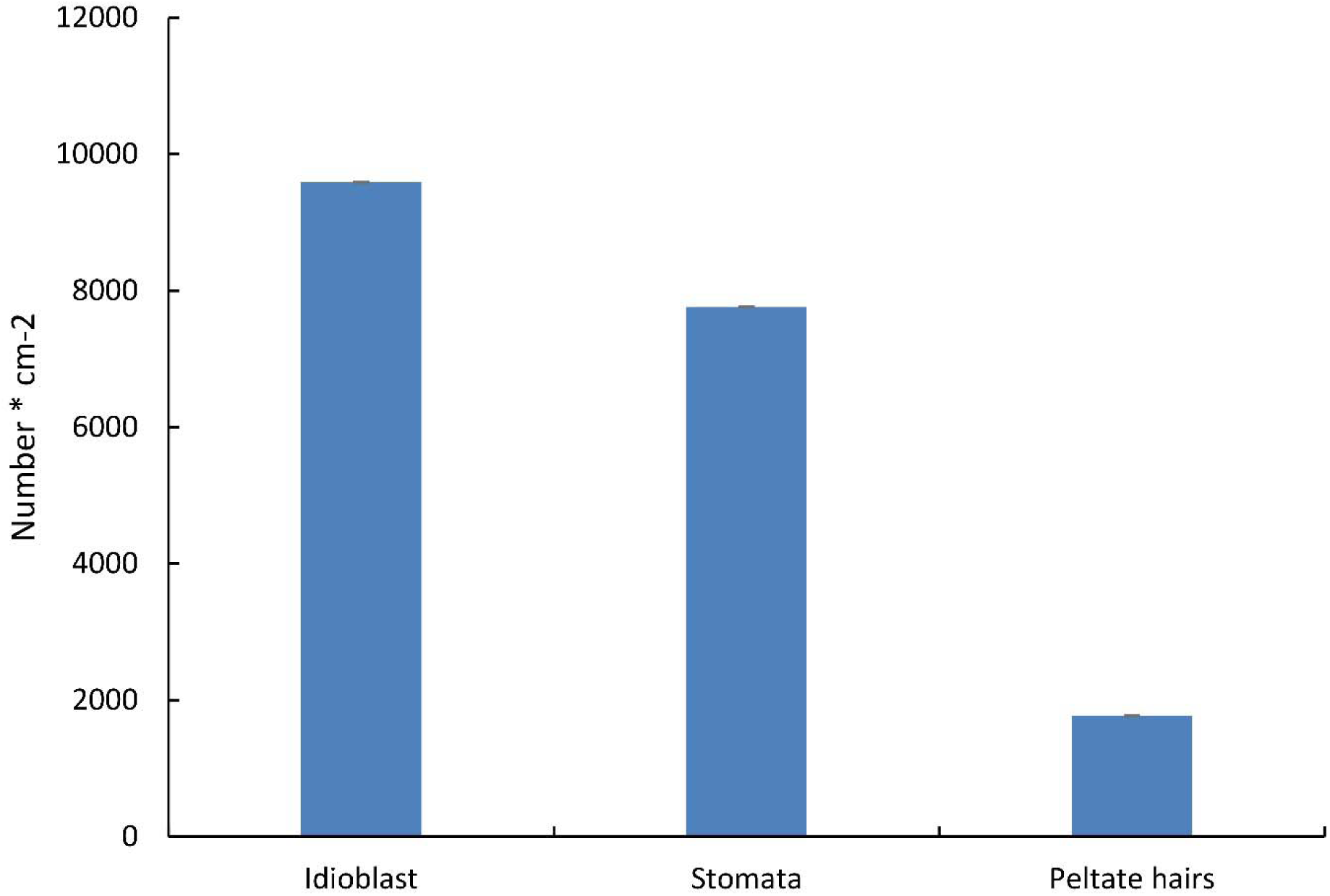
Frequency of idioblasts, stomata and peltate hairs in the leaves of *Capparis odoratissima*. Idioblasts constitute a significant part of the leaf structures, being more numerous than stomata. Interestingly, there are on average ten times more peltate hairs than idioblasts.

### Leaf water uptake in *Capparis odoratissima*

The initial water content of detached leaves was approximately 10 mg cm^−2^. Weight loss of leaves exposed to room temperature conditions was used to understand cuticle evapotranspiration (Fig. 5A). The percentage of water retained by leaves was higher than 80% during the first 21 hours of exposure, but then decreased sharply, being completely dry 143 hours after the initial measurements. In contrast, the cumulative water uptake by submerged leaves was sixty orders of magnitude higher than water losses during the first 10 hours after immersion, surpassing cuticle evapotranspiration even though the cycles for dry and wetting periods were similar (Fig. 5B). Given the asymmetric anatomies between both leaf surfaces, the contribution of each side to water uptake was evaluated by loading them individually with water droplets. As expected, this assay evidenced that water uptake from droplets deposited on only one leaf surface was slower than from submerged leaves, given the effect of cuticle evapotranspiration (Fig. 5C). Additionally, while there was a net uptake of water from both sides, the water balance was positive for the leaves loaded on the abaxial surface only the first two hours after exposure, decreasing thereafter. In contrast, weight gain was maintained for at least ten hours when droplets were applied on the adaxial surface. Most strikingly, the slope of the absorption curve on the adaxial side was the same as the water lost by cuticle transpiration, suggesting that the adaxial surface is responsible for most of the cuticle transpiration, or, in other words, cuticle transpiration is almost suppressed at the abaxial surface.

**Figure 5.**
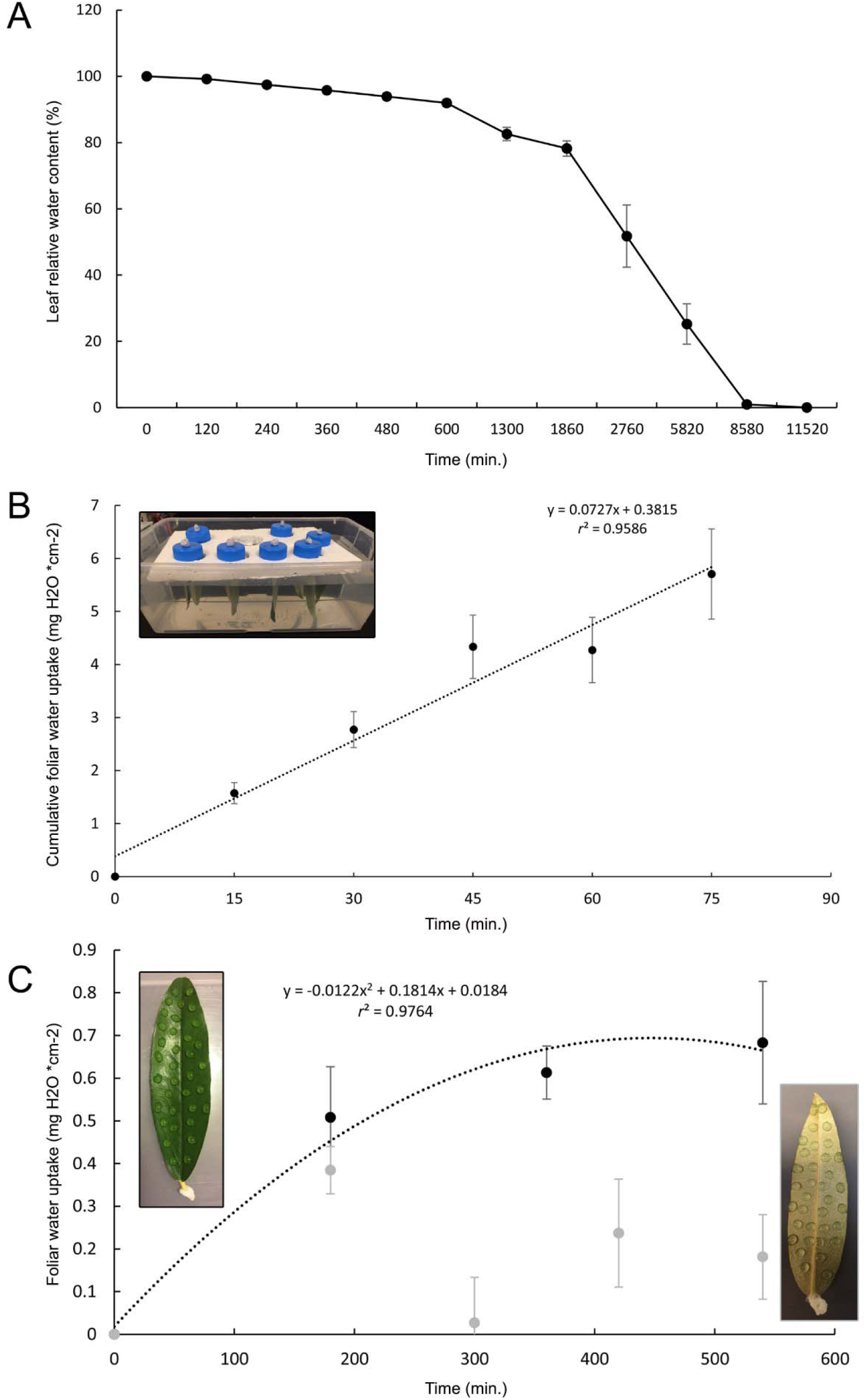
Water uptake by leaves of *Capparis odoratissima*. **A** Relative water content (%) of *C. odoratissima* leaves drying under lab conditions (temperature 22°C; RH 30%), calculated according to van Gardingen and Grace, 1992: RWC_t_=[(Fresh_t_-Dry)/(Fresh_0_-Dry)]*100. **B** Cumulative water uptake by *C. odoratissima* leaves immersed for periodis of 15 minutes followed by quick centrifugation, and then bench dry for 15 min. **C** Foliar water uptake by surfaces of *C. odoratissima* leaves; water droplets on the adaxial surface (black) increased leaf weight in a logarithmic fashion, whereas water droplets in the abaxial surface gained water very modestly (grey dots).

### The pathways of leaf water uptake in *Capparis odoratissima*

Dye loading in the abaxial surface resulted in uptake mediated by the central anchoring area of the peltate hairs (Fig. 6A), evidenced by the intense staining that remained in the thick walls of this central area of the peltate hairs after dye absorption (Fig. 6B). Fluorescence was further observed in the mesophyll of the leaves when loading the dye in either the adaxial or the abaxial surface, pointing to the dual role of the leaf surfaces in water uptake. Strikingly, the structures which fluoresced the most were the walls of the idioblasts, compared with the areas of the leaf without dye (Fig. 6C, D), suggesting that these walls had some degree of hygroscopicity. Not only the external walls, but the lumen of the idioblasts was loaded with the fluorescent dye, including the crenations (Fig. 6E, F), revealing them as the bridges of water uptake from the surface toward the mesophyll.

**Figure 6.**
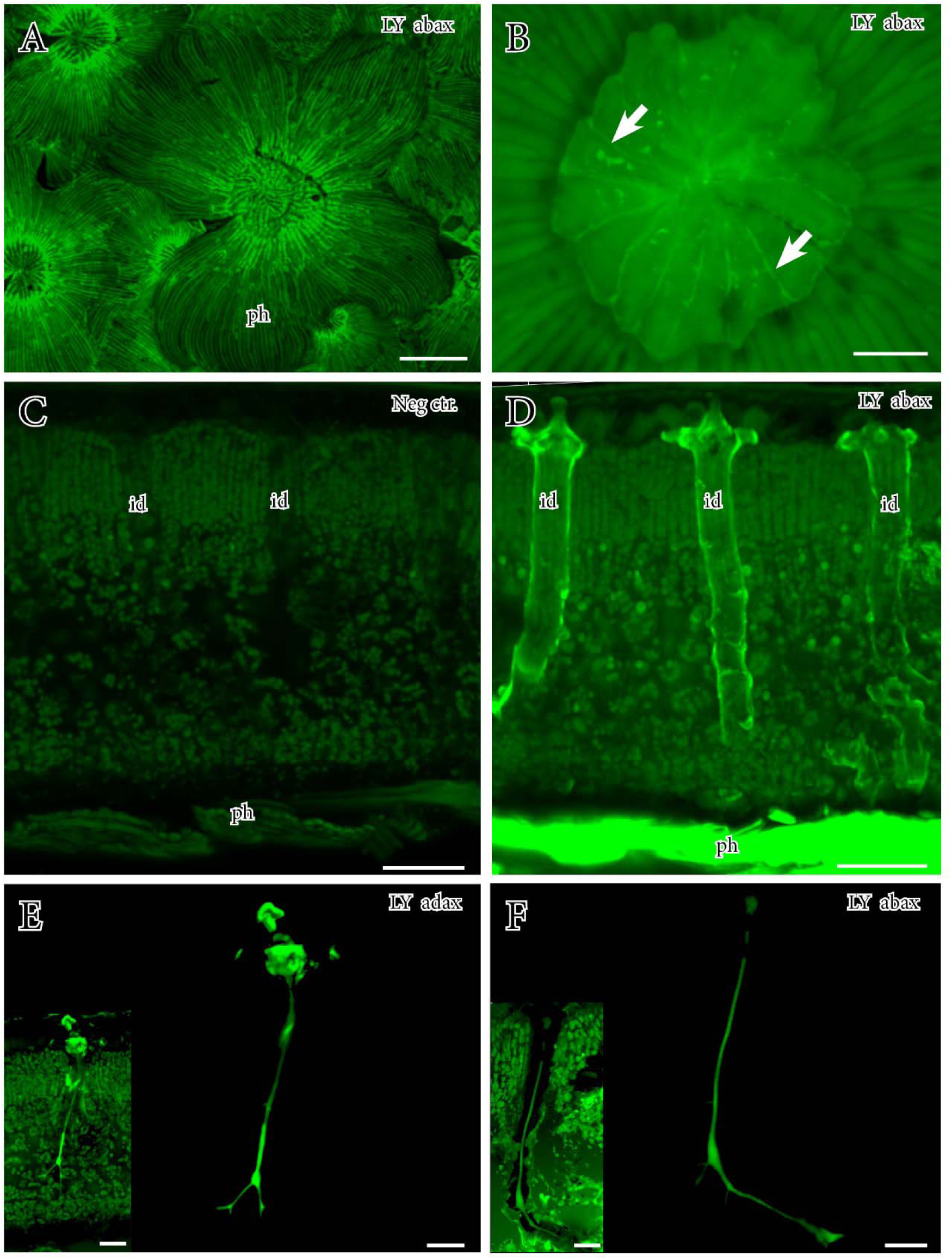
Fluorescent dye loading in the leaves of *Capparis odoratissima*. **A** Negative control showing absence of autofluorescence in the cell walls of the idioblasts with a 488nm wavelength. **D** Following abaxial loading of the dye, idioblasts showed a strong fluorescence on their walls. **E** Maximum projection of the lumen of an idioblast filled with LY dye after adaxial deposition. **F** Same as E, but when the dye was loaded abaxially. **A**. Abaxial deposition of 1µL droplets LY revealed strong fluorescence towards the center of the peltate hairs. **B** Detail of the small external part of the peltate hairs showing fluorescence in the cell walls (arrows). Id, idioblast. LY addax, adaxial load of Lucifer yellow; LY abax, abaxial load of Lucifer Yellow. Scale bars: A = 100µm; B-F = 20µm.

### Pectins and arabinogalactan proteins in the leaves of *Capparis odoratissima*

Immunolocalization of pectin and arabinogalactan protein epitopes with monoclonal antibodies showed a dual topological pattern. On one hand, pectin epitopes were present in the cell walls of both adaxial and abaxial epidermis (Fig. 7A, B), in the external walls of the cells composing the spongy mesophyll (Fig. 7C), and the external cell walls of the peltate hairs (Fig. 7D), but not in the idioblasts.

**Figure 7.**
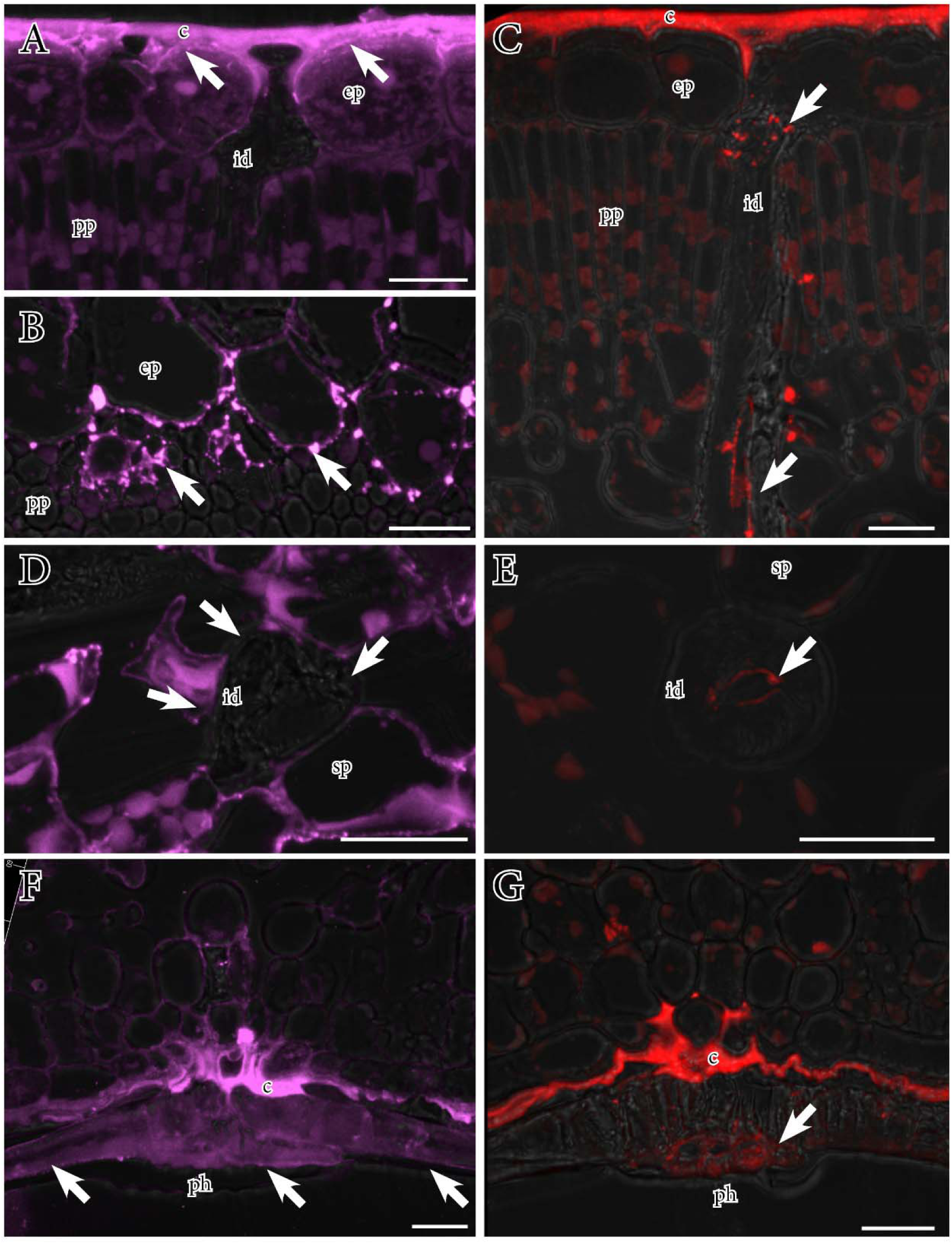
Immunolocalization of pectins (JIM7 mAb) and arabinogalactan proteins (JIM8 mAb) in *Capparis odoratissima* leaves. A, C, F, G, cross-setions of the leaves. B, D, E. Paradermal sections of the leaves. **A, B** Pectins were strongly present in the apoplast of the epidermal cells, particularly between the cuticle and the cell walls of the epidermis (arrows). **C** In contrast, AGPs epitopes were present in the lumen of all the idioblasts (arrows). **D** Pectins also abounded in the external cell walls of the spongy mesophyll in contact with the idioblasts (arrows). **E** Arabinogalactan protein epitopes showed a consistent presence along the lumen of idioblasts (arrow). **F** The external walls of the peltate hairs showed a consistent presence of pectins (arrows). **G** The lumens of the peltate hairs further contained AGPs, consistent with their presence along the idioblast-hair continuum. C, cuticle; ep, epidermis; pp, palisade parenchyma; id, idioblast; sp, spongy parenchyma; ph, peltate hair.

On the other hand, the presence of arabinogalactan proteins was roughly detected with the beta glucosyl Yariv Reagent, which showed an intense red color at the areas of the leaves with idioblasts. More specifically, the epitopes recognized by the JIM8 monoclonal antibody, labelled a narrow area that corresponded with the lumen of the idioblasts, from their adaxial projection (Fig. 7E), through the columnar lumen in the palisade parenchyma (Fig. 7F), to the lumens of the anchoring areas of the peltate hairs (Fig. 7G). As a result, AGPs coated the continuum of lumens between the peltate hairs and idioblasts.

## DISCUSSION

### Water balance and leaf permeability in *Capparis odoratissima*

Previous studies of *C. odoratissima* showed that, unlike other species coexisting in the same environment, *C. odoratissima* increased its biomass with the only source of water being from the atmosphere (Díaz and Granadillo, 2005). However, the magnitude of water uptake by individual leaves is missing so far. Our detailed evaluations of water evaporation through time (i.e. cuticle evapotranspiration), and entry of water (measured as the increase of leaf weight) from detached leaves served to understand the extreme values of water inputs and outputs. First, we demonstrate a high capacitance of the *C. odoratissima* leaves, which kept a high water content (80%) when left to dry with the petiole sealed, in line with the idea of low cuticular permeability for species from xerophylous environments compared with temperate species (reviewed by Kerstiens, 1996). For example, *Fagus sylvatica* leaves showed a more continuous dehydration that responded to water pressure deficit (Gardinen and Grace, 1992). The dehydration rate of *C. odoratissima* leaves is likely lower in natural conditions, not only because the leaves are not detached, but also because the values of relative humidity of the atmosphere oscillates between 50% and 90% (Diaz and Granadillo, 2005). To understand whether water uptake is possible with zero cuticular conductance, we exposed the leaves to a water-saturated environment, revealing a linear uptake of water without cuticle evapotranspiration, gaining up to 40% of the initial water content of the leaves. Leaf water absorption has been widely studied in the major groups of angiosperms, including magnoliids (8 genera), monocots (7 genera), and eudicots (67 genera) (reviewed by Berry et al., 2019 and Dawson and Goldsmith, 2018), pointing to FWU as a key factor affecting plant function in most ecosystems (Weathers et al., 2019). However, this effect is stronger in dry or semi-dry environments (Schreel et al., 2019), such as the dryland tropical areas where *C. odoratissima* grows, where the water in the soil is a limiting resource, and aerial water becomes pivotal for plant growth and survival.

The major barriers for water absorption and/or evaporation are the leaf coatings, such as cuticles. Our anatomical evaluations reveal two different cuticle layers in the two surfaces of *C. odoratissima* leaves. On the upper surface, a thick cuticle layer becomes thinner at some concave areas. Thick lipid coatings such as cuticles associate with avoidance of leaf dehydration (Shields, 1950). Yet, the leaf cuticle is gradually revealing as a permeable layer to absorb water and electrolytes (Fernández et al., 2014; 2017). However, cuticle thickness has not a direct relationship with impermeability (Schuster et al., 2017), but its biochemistry appears to be more important for water exchange between the mesophyll and the atmosphere. The compact and rugose appearance of the cuticle in the lower side of the leaves of *C. odoratissima* suggests a higher surface area and probably a better isolation of leaf tissues. Stomata embedded in this cuticle are gates of water escape through transpiration, but they are covered by a carpet of peltate hairs, which dramatically reduce evapotranspiration. Pubescence is a typical character of xerophylous species, and functionally related with protection against desiccation and thus temperature regulation (Shields, 1950; Fahn, 1986).

### Asymmetric foliar water uptake and anatomical specializations in *Capparis odoratissima*

Our experiments with water droplets in each leaf surface revealed an asymmetry regarding the functional role of leaf surfaces in water uptake. While both surfaces initially showed a positive gain of water, only the adaxial surface maintained this competence longer in time. Water condenses most likely on the adaxial surface of leaves, and, indeed, most works spraying water on the leaf surfaces evidenced that the upper side is more permeable to water (Gardingen and Grace, 1992; Fernández et al., 2014). In *C. odoratissima*, the reasons behind this asymmetric behavior correlate with the markedly distinct anatomy between leaf surfaces. Abaxially, trichomes are hygroscopic, participating in foliar water uptake. Hygroscopic peltate hairs have been reported in some angiosperm species (Gramatikopoulos and Manetas, 1994; Bickford, 2016; Eller et al., 2016; Pina et al., 2016; Vitarelli et al., 2016), being most outstanding in epiphytic bromeliads (Benzing and Burt, 1970; Benzing, 1976; Benzing et al., 1978; Benz and Martin, 2006; Ohrui et al., 2007). However, the presence of peltate hairs is not indicative of foliar water uptake (Bickford, 2016), since remarkable examples with peltate hairs such as in *Olea europaea*, showed no evidence of FWU (Arzeee, 1953). The ability to uptake water by peltate hairs is offset by cuticle evapotranspiration of the adaxial surface (not loaded with water droplets). Thus, a tradeoff between cuticle transpiration and water entrance from the atmosphere appears to be rather positive when water is condensed on the adaxial surface. Oddly, our observations revealed unique and numerous microscopic openings projecting to the atmosphere in the upper leaf surfaces of *C. odoratissima*. Natural openings (i.e. not created by microorganisms) are unusual in this surface, except for amphistomous leaves, or leaves with hydathodes (Martin and von Willert, 2000). We hereby expose unique micropores projecting toward concave areas on the adaxial leaf surfaces, thus allowing symplasmic water transport of water from the atmosphere to the leaf. These openings connect with the lumen of columnar idioblasts, pointing to these structures as significant players in the water budgets of the leaves. In addition, the lumen of the idioblasts formed a continuum with the lumen of the peltate hairs, traversing the cross sectional area of the leaves. As a result, an intricate network of micro channels linked the surface with the mesophyll. Trichome-idioblast associations are commonly found in species from arid environments such as those from the family Euphorbiaceae (Solereder, 1908; Metcalfe and Chalk, 1950), *Olea europaea* (Arzeee, 1953), or *Androstachys johnsoni* (Alvin, 1987). Idioblasts have rarely been demonstrated as vectors of foliar water uptake in angiosperms, with some exceptions such as *Hakea suaveolens* (Heide-Jorgensen, 1990). However, their topology in the numerous forest species where they have been described - including gymnosperms (Hooker, 1864; Sterling, 1947), and more than eighty eudicot families (Solereder, 1908; Foster, 1955a,b; Rao and Mody, 1961; Zhang et al., 2009; Vitarelli et al., 2016), suggested a role in leaf capacitance. As thick-walled sclerenchymatous tissues (Evert, 2006), idioblasts evolved multiple shapes and dispositions within leaves, but the columnar type of idioblasts displayed in the leaves of *C. odortatissima* have only been described in a couple of species so far: *Hakea suaveolens* (Heide-Jorgensen, 1990), and *Mouriria huberi* (Foster, 1947). These adaptations cannot be related to xeric environments, since other species from the same genus and adapted to dry climates have completely different anatomies, such as the Mediterranean *Capparis spinosa* (Rhizopoulou, 1990; Rhizopoulou and Psaras, 2003; Gan et al., 2013).

### The biochemistry of foliar water uptake in *Capparis odoratissima*

The pathway of atmospheric water entry in the leaves of *C. odoratissima* involves hygroscopic materials deposited in the leaf coatings (Gouvra and Gamatipoulus, 2003). In the current work, we first describe the thick walled structures composing the multicellular peltate hairs, which project pectins to the external part, suggesting their involvement in the initial water capture when condensation happens in the abaxial side. A few reports have revealed the presence of pectins in trichomes, such as species from semi-arid forests like *Crombretum leprosum* (Pina et al., 2016), or the tropical species *Drymis brasiliensis* (Eller et al., 2013). In addition, we revealed that the cell walls of both epidermal cells and spongy mesophyll cells show a high concentration of low sterified pectins in their cell walls. This is in line with the ubiquity of pectins within the leaf mesophyll, typically acting as a pulling force for water uptake from the soil. In the leaves of *C. odoratissima*, their tight association with the numerous idioblasts is likely acting as a pulling force for water deposited on the leaf surfaces, using idioblasts as carriers.

The idioblasts of *C. odoratissima* are highly hygroscopic, with cellulosic walls that contain polar molecules for water attachment, but also partially lignified, suggesting a secondary role on defense and structural support. A finely tuned water uptake in *C. odoratissima* is revealed by our experiments with the apoplastic dye tracer Lucifer Yellow. Water entering from the surrounding atmosphere to the idioblasts flows through an intricate network of crenations and branches within the idioblasts that connect all leaf tissues and the exterior. Most strikingly is the fact that epitopes belonging to arabinogalacatan proteins are specifically located within these channels. AGPs are highly branched proteins with a short aminoacid backbone attached to the plasma membrane of cells, and a large saccharidic part that has been related with nutritive and/or signaling functions between cells (Ellis et al., 2010). AGPs play different roles in plant development, including mate recognition and support during reproduction, proper early seedling development, and so many others (Majewska-Sawka and Nothnagel, 2000; Vaughn et al., 2007; Pereira et al., 2016). However, the role of AGPs on plant hydration has been scarcely studied. Remarkably, works evaluating the compositon of the cell walls in the resurrection plant *Craterostigma wilmsii*, which completely losses water, regaining it thereafter, pointed to the hygroscopic properties of AGPs as critical players in this rapid and effective rehydration (Vicré et al, 2004). AGP-related proteins have been previously related with the tensile strength of stems due to their participation in secondary cell wall composition, such as in the vessels or fibers (Ito et al., 2005; Liu et al., 2013). But the presence of AGPs have never been reported before in idioblasts with such specific pattern as in the current work. What it reveals is the biochemical complexity of sclerenchymatous tissues, which are possible effectors enabling hydration of tissues during periods where water deficits in the soil combines with water saturation in the atmosphere (Fig. 8). Thus, AGPs may be secluded in the lumen of the idioblasts during development, serving as bridges of water uptake between the atmosphere and the leaf tissues. Future works will elucidate the possible analogy between AGPs in other sclerenchymatous tissues of other species, as well as their putative role on leaf capacitance.

**Figure 8.**
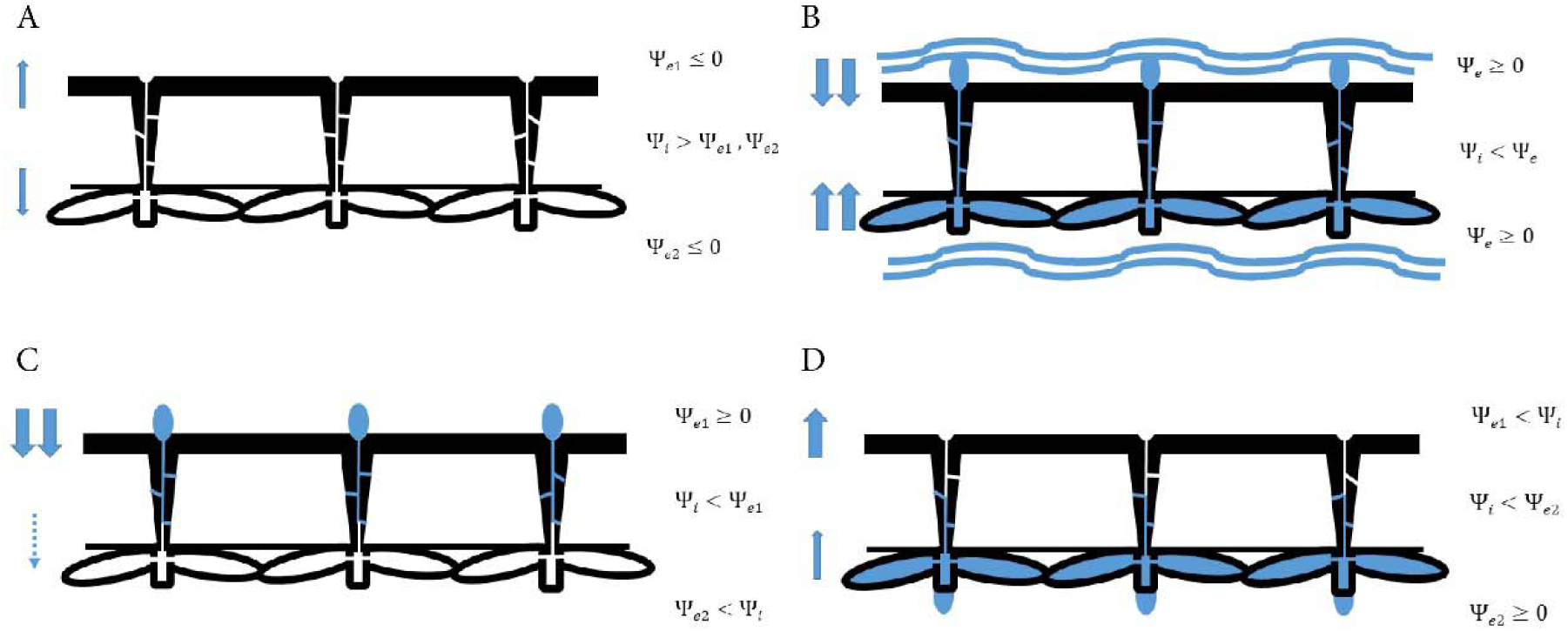
Simplified model of water balances in the leaves of *Capparis odoratissima*. **A.** In the absence of high humidity of the surrounding atmosphere, the water potential of the leaves is higher than the surrounding environment, thus there is a net loss of water (blue arrows). **B**. The opposite situation happens with submerged leaves, which have lower water potentials than the environment, thus gaining water (blue arrows) through the lumen of the continuum peltate hairs-idoblasts. **C**. When water condenses in the upper surface, the inner leaf potential is lower, thus allowing water entrance through he lumens of the idioblasts (blue arrow), and allowing a minimal evapotranspiration in the lower surface due to the presence of an extra layer of peltate hairs covering the stomata (dotted arrow). **D**. When water condenses in the lower surface, water enters through the lumen of the peltate hairs and the idioblasts (thin blue arrow), but evapotranspiration of the upper surface appears to be higher than water entrance (thick blue arrow).

## CONCLUSIONS

Our data strongly suggest that the pathway for water uptake in *Capparis odoratissima* is mediated by the idioblasts and peltate hairs, which are interconnected. We propose a unique model in angiosperms (Fig. 8) that involves apertures of the leaves toward the adaxial surface, which are involved in water uptake, but may facilitate evapotranspiration. Hygroscopic materials belonging to arabinogalactan proteins serve to capture water to the lumen of the idioblasts, and pectins in the mesophyll and the epidermis may further facilitate the incorportation of water to the leaf tissues. This cascade of biochemical and physical events allows *Capparis odoratissima* trees to make use of atmospheric water resources.

## ACKNOWLEDGEMENTS

We are grateful to Ned Friedman, Kea Wodruff, and Faye Rosin from the Arnold Arboretum of Harvard University for sharing facilities. We thank Mr. Omar Fernando Sierra for his help with plant material. Juan M. Losada is a ComFuturo Researcher at the IHSM-CSIC-UMA, funded by the Fundación General CSIC (FGCSIC), and within the project RTI2018-102222A100. This work was supported by the National Science Foundation (EAGER 1659918, IOS 1456845 and DMR 14-20570).

